# Tuning electrical stimulation of the cervical vagus nerve for abdominal signaling while reducing cardiovascular side effects

**DOI:** 10.1101/2020.12.10.420398

**Authors:** Charles C. Horn, Mats Forssell, Michael Sciullo, Jonathan E. Harms, Stephanie Fulton, Chenchen Mou, Fan Sun, Tyler W. Simpson, Gutian Xiao, Lee E. Fisher, Christopher Bettinger, Gary K. Fedder

**Affiliations:** Department of Medicine, University of Pittsburgh, Pittsburgh, PA, USA; UPMC Hillman Cancer Center, Pittsburgh, PA, USA; Center for Neuroscience, University of Pittsburgh, Pittsburgh, PA, USA; Department of Electrical and Computer Engineering, Carnegie Mellon University, Pittsburgh, PA, USA; Department of Biomedical Engineering, Carnegie Mellon University, Pittsburgh, PA, USA; Department of Microbiology and Molecular Genetics, University of Pittsburgh, Pittsburgh, PA, USA; Department of Physical Medicine and Rehabilitation, University of Pittsburgh, Pittsburgh, USA; Department of Bioengineering, University of Pittsburgh, USA; Department of Material Science and Engineering, Carnegie Mellon University, Pittsburgh, USA

**Keywords:** vagus nerve, vagus nerve stimulation, bioelectronic medicine, electroceuticals, neuromodulation

## Abstract

Electrical vagus nerve stimulation (VNS) has the potential to treat a wide variety of diseases by modulating afferent and efferent communication to the heart, lungs, esophagus, stomach, and intestines. Although distal vagal nerve branches, close to end organs, could provide a selective therapeutic approach, these locations are often surgically inaccessible. In contrast, the cervical vagus nerve has been targeted for decades using surgically implantable cuff electrodes to treat epileptic seizures and depression; however, to date, clinical implementation of VNS has relied on an electrode with contacts that fully wrap around the nerve, producing non-selective activation of the entire nerve. Here we demonstrate selective cervical VNS using cuff electrodes with multiple contacts around the nerve circumference to target different functional pathways. These flexible probes were adjusted to the diameter of the nerve using an adhesive hydrogel wrap to create a robust electrode interface. Our approach was verified in a rat model by demonstrating that cervical VNS produces neural activity in the abdominal vagus nerve while limiting effects on the cardiovascular system (i.e., changes in heart rate or blood pressure). This study demonstrates the potential for selective cervical VNS as a therapeutic approach for modulating distal nerve branches while reducing off target effects. This methodology could potentially be refined to treat gastrointestinal, metabolic, inflammatory, cardiovascular, and respiratory diseases amenable to vagal neuromodulatory control.

## 1. Introduction

Neuroelectric therapeutic devices targeting peripheral nerves hold the promise to effectively treat diseases without the side effects often associated with systemic administration of pharmacotherapies [1,2]. Moreover, devices that target the vagus nerve have the theoretical potential to modulate a broad array of peripheral organs, such as the immune system, gastrointestional tract, pancreas, liver, heart, and lungs, which could provide therapies for obesity, diabetes, heart disease, gastrointestinal disease, and inflammatory disorders. This potential, however, has not been realized because methods have yet to be refined for targeting specific functional groups of vagal nerve fibers.

Current approaches to vagus nerve stimulation (VNS) attempt to achieve efficacy while minimizing side effects by: (1) tuning stimulation parameters to each patient [3,4]; and, (2) placing electrodes close to end organs [5,6]. Clinically-available VNS electrodes, such as the Cyberonics helical cuff design [7,8], have a single contact which substantially limits potential for tuning parameters to achieve selective stimulation while reducing off target effects. Side effects of clinical VNS commonly include hoarseness, cough, dyspnea, pain, paresthesia, nausea, and headache [9]. In addition, placement of electrodes close to end organs can be surgically difficult and risky for patients. In contrast, increasing the number and density of electrode contacts around the vagus nerve appears to provide greater functional selectivity, which has been tested for the cardiovascular system [10].

In the current report we developed a cervical VNS approach using a multi-contact electrode to target abdominal organs. We used microfabrication technology to create small contacts placed circumferentially on the vagus nerve and wrapped in hydrogel to achieve intimate contact between the electrodes and the vagus nerve. In *in vivo* anesthetized rats we recorded nerve responses from the abdominal vagus (CAP, compound action potential), as well as blood pressure (BP) and heart rate (HR), to maximize the CAP response while reducing cardiovascular off-target effects.

## 2. Materials and methods

### 2.1. Subjects

Experiments were conducted on 21 male Sprague-Dawley rats (302 – 452 g; Taconic). Rats were food deprived for 139 ± 28 min (mean ± SD) prior to surgery. Animals were otherwise maintained with *ad libitum* access to food and water and kept in a temperature and humidity controlled vivarium on a 0700:1900 h on/off light cycle. All experiments were approved by the University of Pittsburgh Institutional Animal Care and Use Committee.

### 2.2. Electrode fabrication

The probes consist of a connector region, a cable region and an electrode region (Fig. 1) [11]. The connector region is designed to interface with a zero insertion force (ZIF) connector (FH26 Series, Hirose Electric Co., Tokyo, Japan). The meandering in the cable region results in higher compliance, and increases the robustness of the probe to stretching, bending, and torsion. The electrode region is angled at 45° in order to facilitate positioning of the probe with respect to the nerve. Four electrodes (*W × L*, 25 μm by 700 μm) are distributed between two flaps; the electrodes are oriented longitudinally along the direction of the nerve in order to increase the electrode area in contact with the nerve. Each electrode is connected to two conductors from the connector in a loop configuration to allow for testing probe connectivity. A second pair of flaps wrap in the opposite direction around the nerve to help mechanically stabilize the probe during surgery. Holes in the secondary flaps allow the hydrogel to maintain contact with the nerve through the flap.

**Fig. 1.**
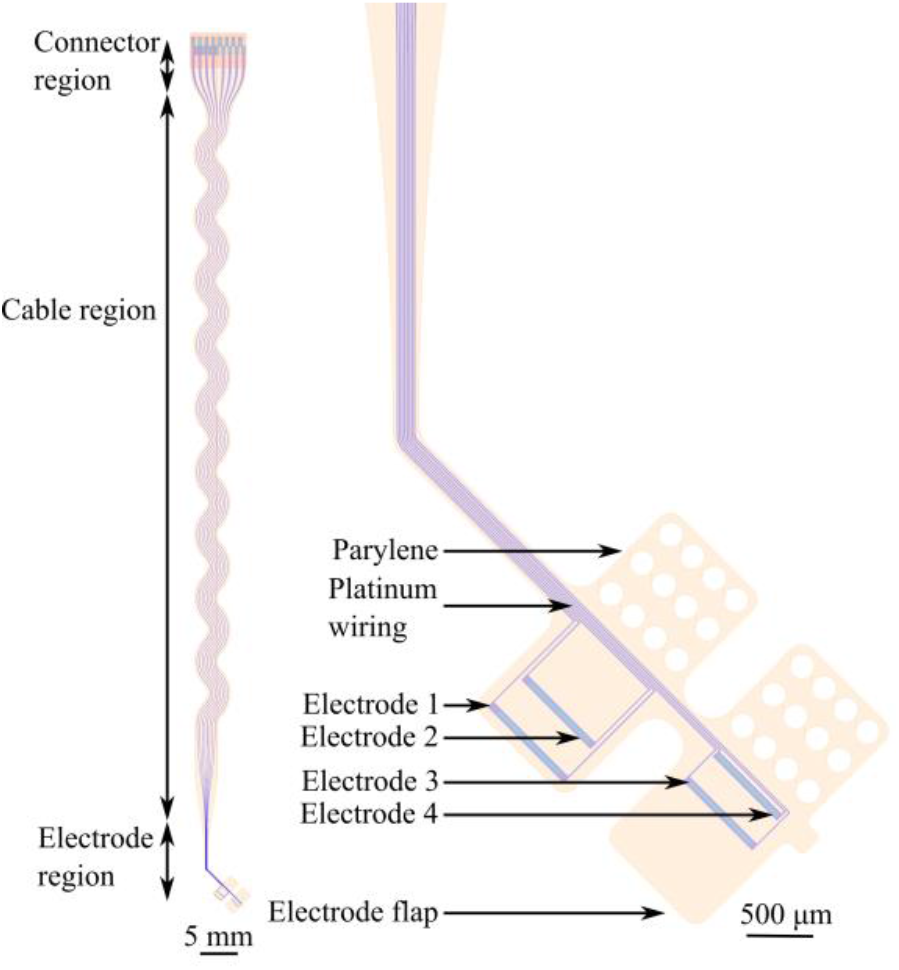
Layout of probe with electrode contacts facing up. Left: full probe; Right: detail of electrode region.

Probes were fabricated on Si handle wafer (100 mm diameter). A sacrificial release layer consisting of polyacrylic acid (PAA) [12] was first spun on the wafer. Then, 2 μm-thick parylene-C was deposited (Labcoter 2, Specialty Coating Systems, Inc., Indianapolis, IN, USA) and holes for electrodes and contacts were patterned using O_2_-based reactive ion etching (RIE; 50 mTorr, 14 sccm O_2_, 50 W; Phantom II RIE, Trion Technology, Inc., Tempe, AZ, USA) and masked with 9 μm-thick photoresist (AZ P4620, Microchemicals GmbH, Ulm, Germany).. The parylene patterning was performed using proximity alignment, which results in a sloped sidewall profile, which allows the recess to be completely covered by the subsequent metal layer, resulting in a flush surface between the metal and parylene. A 120 nm-thick layer of platinum was deposited and patterned using ion milling (Commonwealth Scientific, Wokingham, UK). A second parylene layer identical to the first was then deposited, and the probe periphery is patterned and etched using the O_2_-based RIE.

After dicing the handle wafer, a polyimide-based stiffener (Pyralux LF Coverlay, DuPont, Inc., Wilmington, DE, USA) was attached to the connector region of the probe to allow insertion into the ZIF connector. The diced substrate is oriented at a 30° angle with respect to gravity and with the probe connector region at the bottom. This arrangement allows manual application of polydimethylsiloxane (PDMS, 5:1 base:curing agent, Sylgard 184, Dow Chemical Co., Midland, MI, USA) to flow down from the middle of the cable region. The PDMS allows easier handling of the probe during surgery and improves the robustness of the interface with the ZIF connector. Because the parylene layer is situated at the bottom of the PDMS layer, bending stress can cause the parylene to crack if the PDMS is too thick. The probe is then released and inserted into a ZIF clip connector on a printed-circuit board (PCB). The PCB assembly is coated in PDMS (10:1 base:curing agent, Sylgard 186, Dow Chemical Co., Midland, MI, USA) and placed on a piece of cleanroom paper. The probe is inserted into a plastic straw, lined with paper to eliminate static charging, for manipulation during surgery.

Similar to a prior study [13], the adhesive hydrogel was prepared with 16 mM catechol functionalized four-arm poly(ethylene glycol) [PEG-Cat]_4_ and iron(III) chloride ([Fe3+]:[Cat]= 0.58) in 50 mM Tris base/diH_2_O solution as solvent. Hydrogel disks were 3 to 4 mm in diameter.

### 2.3. Acute surgery and testing

Animals were either anethetized with urethane (ip, initial dose of 1 g/ml/kg and supplemental injections; 4.1 ± 0.7 ml total; N = 8) or isoflurane (placed in a chamber and then transferred to a nose cone, 1 to 3% isoflurane; N = 13). In rats anethesized with isoflurane, we were testing a chronic implant procedure, which included connecting the probe lead to the PCB affixed to the cranium. All procedures were the same for urethane and isoflurane experiments except that, in urethane injected rats, a fluid-filled catheter was inserted into the left carotid artery to measure blood pressure (DBP1000 series Direct Blood Pressure System, Kent Scientific, Torrington, CT, USA), and a tube was placed into the trachea to record respiration (SAR-830/AP Small Animal Ventilator, CWE, Inc., Ardmore, PA, USA). Body temperature was kept at 37 °C by a regulated heating pad. Alligator electrical clips were attached to the right and left flanks to record heart rate (P511 amplifier with low impedance headstage, Grass Technologies). A ~2 cm incision was made in the ventral neck to separate the muscles and fascia from the left vagus nerve. The VNS probe was placed on a disk of hydrogel, with the nerve above, and then the hydrogel was wrapped around the nerve as the opposing edges adhere together (Fig. 2B and 2C). Subsequently, the abdominal cavity was opened with a midline incision, a three-point surgical retractor was placed on the abdominal muscle (left and right) and xiphoid process, the liver positioned to the right side using gauze and a curved spatula retractor, and the stomach was retracted caudally to reveal the ventral vagus on top of the esophagus. A bipolar Pt-Ir wire (50.8 μm diameter; AM Systems, Sequim, WA, USA) electrode was placed around the ventral abdominal vagus nerve and embedded in silicone (Kwik-cast, WPI Inc., Sarasota, FL, USA).

**Fig. 2.**
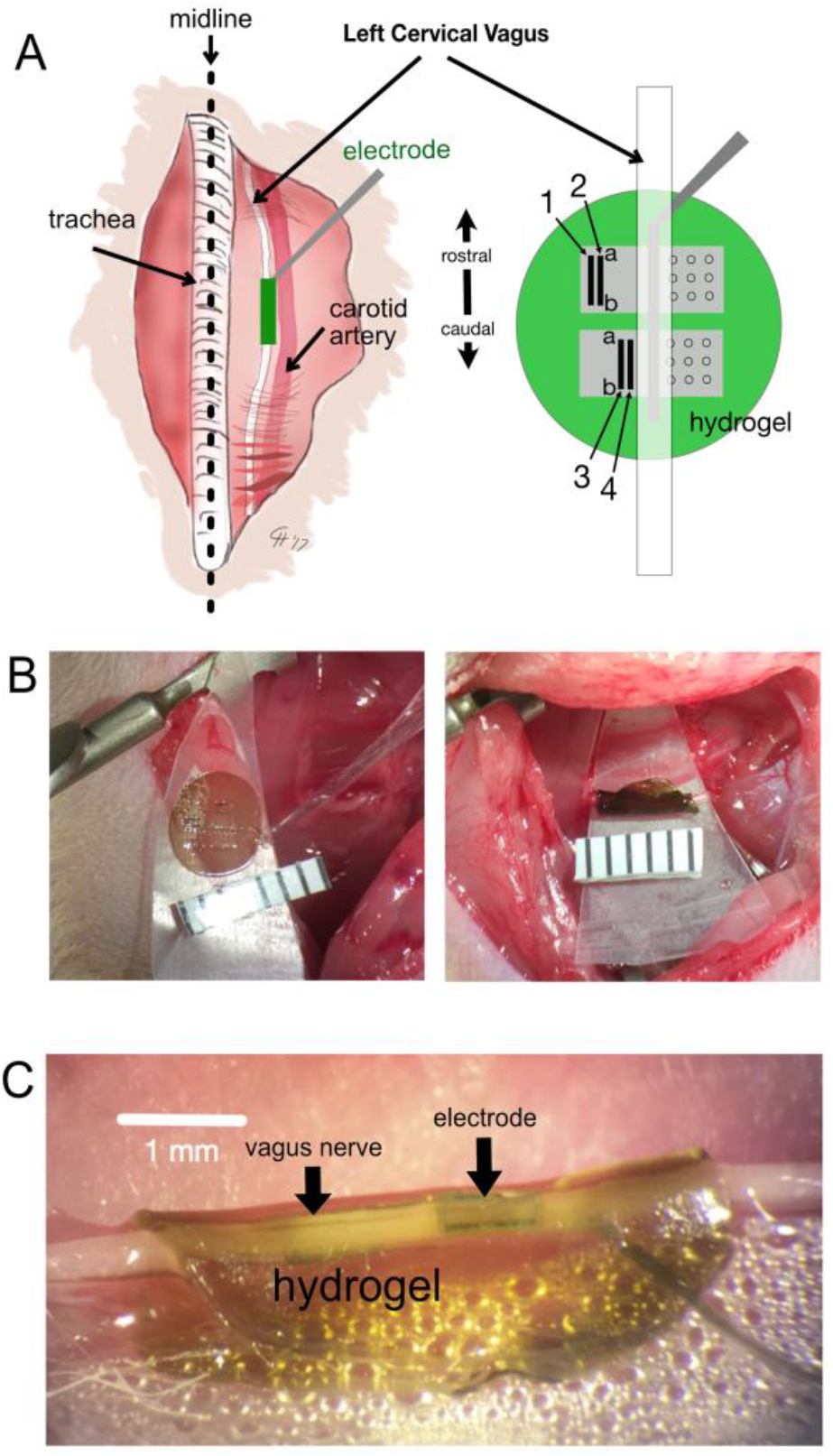
Electrode surgical placement. A) Left cervical vagus nerve: right diagram shows hydrogel under the nerve and probe before wrapping the hydrogel, a and b show rostral and caudal ends of each electrode contact. B) Implantation, before and after hydrogel wrap: ruler has 1 mm scaling. C) High magnification image showing electrode under the hydrogel.

The VNS leads were wired to a custom rail connector with two lead channels (“a” and “b”) for each contact site on the nerve (Fig 1 and 2A); we used these two channels to provide redundancy for each contact in case of breakage and to test continuity across each contact. In total, there were eight channels for stimulation, contacts 1 to 4 and an “a” and “b” connection for each. A separate ground wire was attached to the edge of the surgical incision in the neck using a micro-alligator clip. Impedances were checked (Impedance conditioning module; FHC, Bowdoin, ME, USA) in two ways: (1) between contact redundant pairs, to measure continuity and conductor impedance; and (2) from each of the eight channels to ground, to measure electrode interface impedance. Stimulation was applied using a constant current stimulator (Model 4100; AM Systems, Carlsborg, WA, USA). Abdominal vagus nerve signals were recorded using an AC amplifier equipped with a high impedance headstage (DAM80, WPI Inc., Sarosota, FL, USA). All signals were recorded and stored using a digital I/O interface and software (Power 1401 and Spike2, Ver. 7; CED, Cambridge, England).

After impedances were tested (27.7 ± 26.5, kΩ; mean ± SD; this is the average impedance for those values less than 100 kΩ, which we considered as a broken connection), monopolar stimulation was applied to each channel (0.5 ms pulse duration; 10 pulses for each test, 2 Hz) starting at 1 mA and currents were adjusted up and down (maximum current was 2 mA) across channels to acheive a CAP in the abdominal vagus nerve and decrease the effects on BP and HR. At the end of experiments, animals were euthanized with an overdose of Beuthanasia, intracardiac injection.

### 2.4. Data analysis

Data analysis was conducted by importing Spike2 CED time series data into Python within Jupyter notebooks (Neo; https://github.com/NeuralEnsemble/python-neo). For each animal we computed the effect of VNS on CAP and HR (and in some cases BP). For CAP, to remove the stimulus artifact and sample the dominant C-fiber activity in the vagus nerve, we used a window at 40 to 140 ms post-VNS for the VNS test period and 140 to 240 ms for the baseline period (Fig. 3A): this time window was based on our prior work showing VNS C-fiber responses in the rat vagus nerve [14]. For BP, we used the 10 s before VNS for the baseline and the 10 s after the start of VNS for the test period (Fig. 3A). For HR, we computed the ECG inter-event intervals (time between each ECG), and used 7 s before VNS and 7 s after the start of VNS (Fig. 3A). We selected these time periods to capture all of the effects of VNS on these physiological measures. To make comparisons between the above three measures (CAP, BP, and HR) we computed Z-scores for the test period compared to baseline (Fig. 3B). The effects of VNS were also computed as an average of 10 stimulation pulses.

**Fig. 3.**
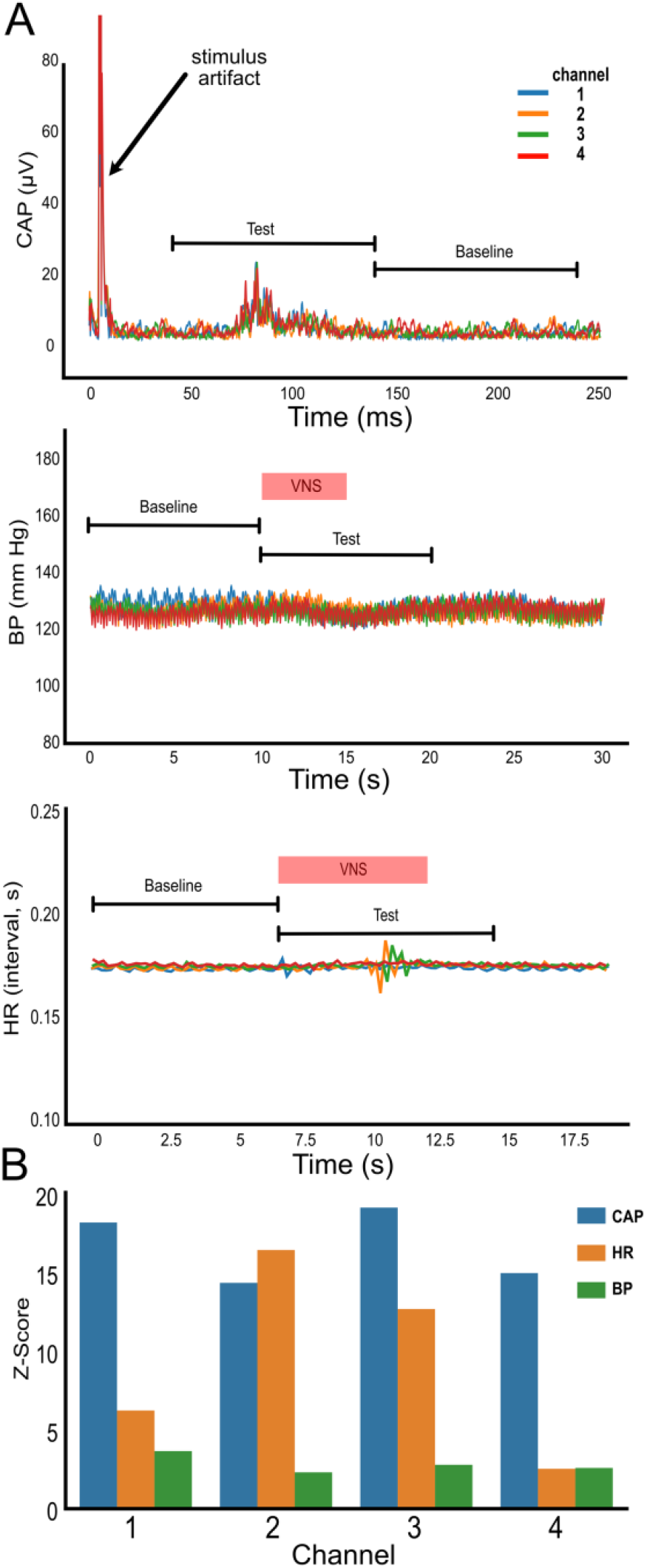
Representative sample of physiological signal processing from one animal; all channels have a CAP response but only channels 1 and 4 selectively for CAP vs. HR. **A)** Signals from CAP, BP, and HR showing the baseline and test periods used to compute the Z-scores. **B)** Comparison of Z-scores for each channel. Channel 4 shows the most selective (“Best”) response of VNS on abdominal CAP.

Ratios of Z-scores were then computed using CAP Z-score divided by BP Z-score (CAP/BP) and CAP Z-score divided by HR Z-score (CAP/HR). For each animal, the “best” (i.e., a high ratio value; more selective for CAP) and “worst” channels (i.e., a low ratio value; less selective for CAP) were selected for each of these ratios. Data were analyzed to determine their fit to a normal distribution using the “normaltest” function in the Scipy package of Python (https://docs.scipy.org/doc/scipy/reference/generated/scipy.stats.normaltest.html). This is an omnibus test of skew and kurtosis. This test indicated the data were not from a normal distribution for the CAP/HR ratio; therefore, we chose to use non-parametric Mann-Whitney U tests to compare groups, with p < 0.05 as statistically significant.

Finally, we produced visualization of the scatterplots of Z-scores for CAP vs. BP, CAP vs. HR, and BP vs. HR, including a linear regression fit (Python seaborn package; https://seaborn.pydata.org/). We also computed the Pearson R correlation coefficients for these relationships and used p < 0.05 as a criterion for statistical significance (Scipy, Python statsmodels package). Our correlation analyses revealed that outlier values substantially skewed the results toward statistical significance, e.g., the analysis of BP versus HR; therefore, we trimmed the distributions by removing values that were > 3 standard deviations from the mean to create more robust statistics (this removed three values from the analysis).

## 3. Results

Twenty-two percent of the redundant traces showed an unacceptable high impedance level (> 100 kΩ); however, because of this redundancy most subjects had four working channels (18 of 22 animals), in contrast to those with three working channels (2 out of 22 animals) and only one working channel (2 out of 22 animals).

The ratios of best and worst channels for CAP selectivity compared to cardiovascular responses were statistically significant using Mann-Whitney U tests (CAP/BP, U=7.0, p=0.005; CAP/HR, U=16.0, p=0.001; Fig. 4). Linear fitting of CAP to BP or CAP to HR indicated no consistant relationships (Fig. 5), and the correlation coefficients were also not statistically signicant (CAP vs. BP, r = 0.007, p = 0.97; CAP vs. HR, r = −0.007, p = 0.58; BP vs. HR, r = 0.36, p = 0.07).

**Fig. 4.**
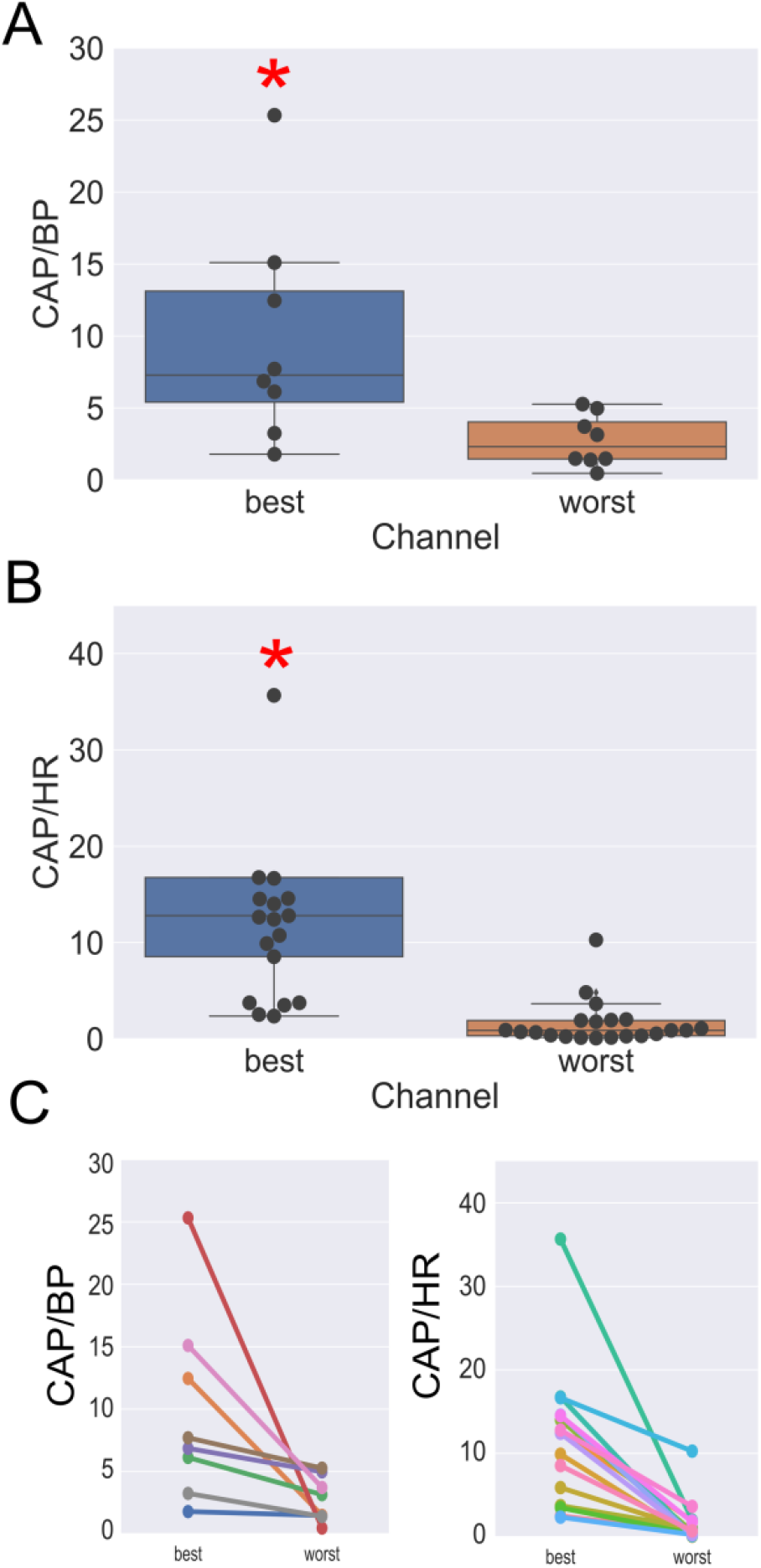
Plots showing “best” and “worst” nerve cuff channels for selectivity of CAP compared to cardiovascular responses. **A)** CAP/BP ratio (compound action potential to blood pressure) and **B)** CAP/HR (CAP to heart rate). Best channel is the highest ratio and worst channel is the lowest ratio. *, p < 0.05, Mann-Whitney U test. Note that for Fig. B, three values in the “best” condition are out of range for the y-axis (323, 128 and 2461). C) Line plot with different colors showing “best” and “worst” channels for each animal.

**Fig. 5.**
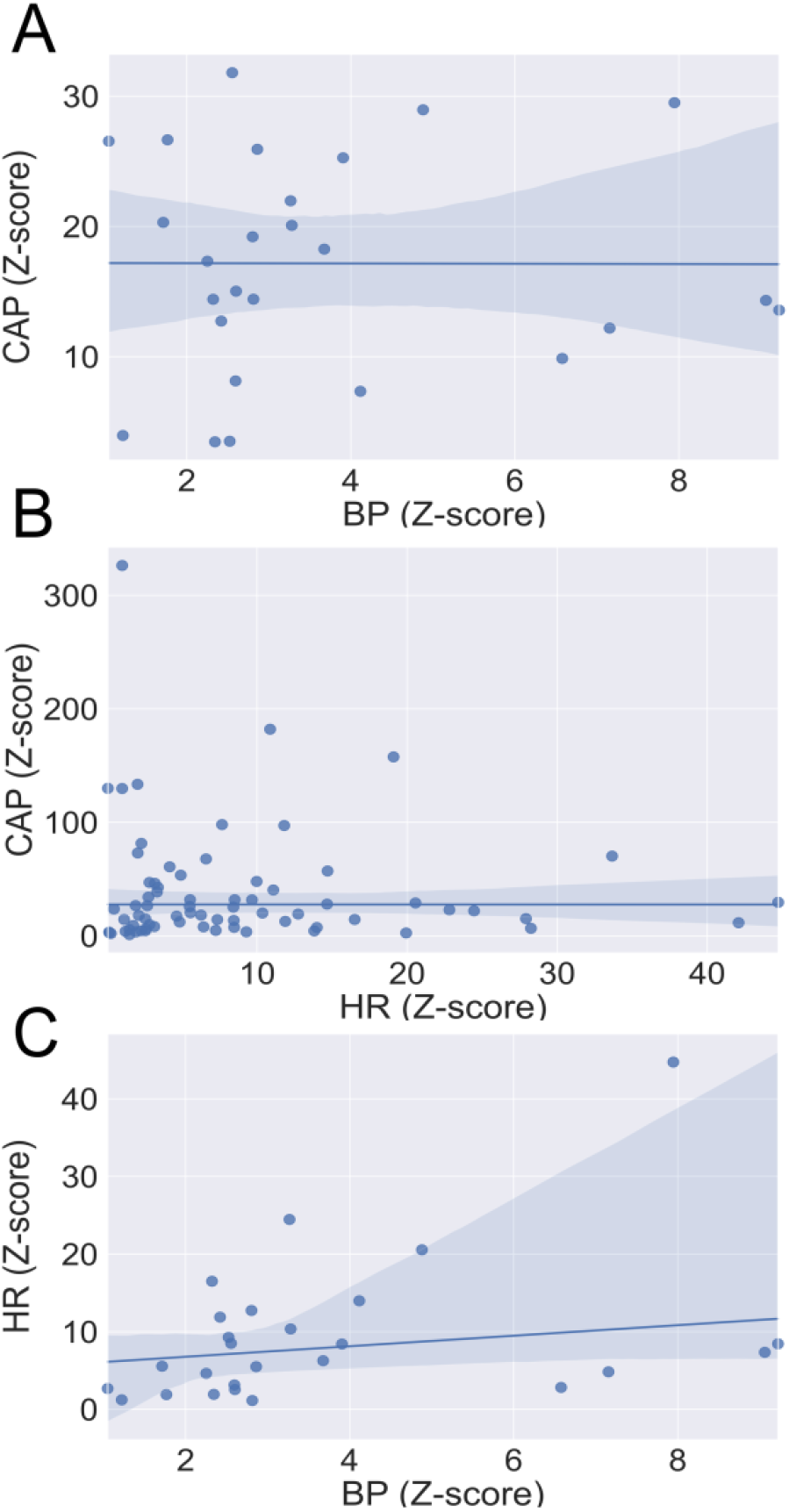
Scatterplots and linear regression model fit (line) of Z-scores showing the relationships for **A)** CAP versus BP (compound action potential vs. blood pressure), **B)** CAP vs. HR (CAP vs. heart rate), and **C)** HR vs. BP. Blue shading indicates the 95% confidence interval of the linear regression fit.

## 4. Discussion

We verified the functional selectivity of multi-contact stimulation of the vagus nerve with a custom microfabricated cuff electrode. Specifically, we showed spatial tuning of VNS to activate abdominal vagal fibers compared to cardiovascular responses. We also demonstrated that CAP and BP (or HR) are not functionally related using regression and correlation analyses.

In addition to functional selectivity, the current approach used a non-traumatic electrode. Unlike traditional nerve cuff electrodes, the current method uses flexible hydrogel, which can ensure a close position and tight seal between the electrode contact and the nerve. The flexible probe along with the adhesive hydrogel allows sufficient electrode-nerve contact to stimulate the cervical vagus, while minimizing nerve compression. Moreover, unlike a traditional cuff electrode, the current hydrogel approach does not rotate; and therefore, the placement of electrode contacts relative to potential functional pathways remains the same.

This modular microfabricated probe platform can easily be adapted to other animal models or probe locations. The electrode count is limited here by the small dimension of the rat vagus (less than 500 μm in diameter) and can be increased for larger nerves, which could allow even greater control of selectivity. Additionally, recording electrodes can also be used to enable closed-loop control of the stimulation parameters.

Future applications of this approach include chronic testing, with assessment of the effects on disease mechanisms. It will also be important to determine how our CAP results relate to activation of specific abdominal organs, as well as understand the role of afferent and efferent fibers. Therapeutic impacts could be wide-ranging for application to disorders associated with the abdominal cavity to include effects on obesity, diabetes, gastrointestinal diseases, and inflammatory disorders.

## Acknowledgements

The authors gratefully acknowledge the staff of the Carnegie Mellon Nanofabrication Facility, and the prior design and fabrication work on electrodes by Dr. Xiao Chuan Ong. We also thank the excellent animal care provided the University of Pittsburgh DLAR at the UPMC Hillman Cancer Center. This work was supported by funding from the Pittsburgh Health Data Alliance.

